# Honey bees cannot sense harmful concentrations of metal pollutants in food

**DOI:** 10.1101/2021.06.15.448345

**Authors:** Coline Monchanin, Maria Gabriela de Brito Sanchez, Loreleï Lecouvreur, Océane Boidard, Grégoire Méry, Jérôme Silvestre, Gaël Le Roux, David Baqué, Arnaud Elger, Andrew B. Barron, Mathieu Lihoreau, Jean-Marc Devaud

## Abstract

Whether animals can actively avoid food contaminated with harmful compounds through taste is key to understand their ecotoxicological risks. Here, we investigated the ability of honey bees to perceive and avoid food resources contaminated with common metal pollutants known to impair their cognition at low concentrations (lead, zinc and arsenic). In behavioural assays, bees did not discriminate food contaminated with field-realistic concentrations of these metals. Bees only reduced their food consumption and displayed aversive behaviours at very high, unrealistic concentrations of lead and zinc that they perceived through their antennae and proboscis. Electrophysiological analyses confirmed that high concentrations of the three metals in sucrose solution induced a reduced neural response to sucrose in their antennae. Our results thus show that honey bees can avoid metal pollutants in their food, but only at very high concentrations above regulatory levels. Their inability to detect lower, yet harmful, concentrations in a field-realistic range suggests that metal pollution is a major threat for pollinators.

## 1. Introduction

Pollinators play major economic and ecological roles by facilitating the reproduction of flowering plants. Worryingly, pollinating insects are declining due to many stressors derived from human activities, among which are pesticides, reduced floral diversity, pests and viruses [1]. Exposure to potentially harmful metal pollutants may have additional impact, though largely overlooked despite raising ecological and public health concern worldwide [2]. The release of metal pollutants into the environment, as a result of industrial manufacturing and mineral extraction, has resulted in their accumulation in ecosystems at levels far beyond concentrations that would be considered natural [3,4]. Because metallic compounds cannot be degraded and can be poisonous at low levels, they represent a potential threat to animals exploiting contaminated resources [2].

In the case of pollinators, such as bees, the effects of metal pollutants could have ecosystemic consequences [2]. Bees are exposed to metal pollutants while flying [5] and collecting food resources (water, pollen and nectar) [6,7]. Metals then bio-accumulate in the bodies of the bees [8,9], as well as in hive products [10,11]. Many deleterious effects of metal pollutants, which vary depending on doses and durations (i.e. chronic vs. acute) of exposure, have been described in mammals [12], birds [13], and specifically on human health [14], and there is clear evidence that exposure to metals can have deleterious effects on the survival [15], physiology [16,17] and behaviour [18,19] of bees. However, whether bees can detect metal pollutants in food is not known.

Bees can detect natural deterrent substances produced by plants and recognize them as harmful, at least in specific experimental conditions [20–23]. Even when ingested, such substances trigger subsequent aversive responses due to a delayed malaise-like state [20,22– 24]. If endowed with sensitivity to harmful metal concentrations, bees could actively avoid contaminated food. They could also consume food containing low concentrations, that are harmless or even profitable, as some metallic compounds are micronutrients needed for physiological functions [25].

Metal ions can distort the function of peripheral chemoreceptors involved in taste-mediated feeding behaviour [26], particularly by reducing the sensitivity of gustatory neurons to sugars in some insect species [27,28]. Honey bees can recognize a variety of potentially noxious substances through gustatory receptor cells located on their antennae, mouthparts and forelegs [29], but their capacity to detect and/or avoid metals in food seems limited. In a study of the proboscis extension reflex, restrained bees willingly consumed solutions containing field-realistic levels of selenium [30] or cadmium [18] with no behavioural indications of avoidance. Copper and lead solutions appeared to be palatable at certain concentrations, and only lead solutions induced any aversive responses [18]. Field studies have reported either no discrimination between flowers grown in lead-contaminated or uncontaminated soils [31], or increased visitation of zinc- and lead-treated flowers [32]. Thus, it appears that the ability to detect and reject potentially toxic substances varies greatly with their chemical identities and concentrations, the body parts in contact with them (mouthparts, antennae or tarsi), and the experimental or ecological context (harnessed, free-flying individuals). Whether bee taste receptors actually respond to metals has never been tested to our knowledge, so that the mechanisms of metal perception remain unknown [18].

Here, we tested whether bees could detect common metals in food. We focused on salts of zinc (an essential nutrient at low concentrations [33]), as well as of lead and arsenic (two major environmental pollutants [34]). We first assessed whether bees modified their consumption of sucrose solutions containing metal pollutants in choice and no-choice conditions. We then investigated whether bees could detect metal pollutant salts through their antennae and proboscis. Finally, we tested the capacity of gustatory antennal neurons to respond to metal pollutant salts delivered alone or in combination with sucrose.

## 2. Methods

### (a) Bees and metals

We collected honey bees (*Apis mellifera*) from fourteen hives at our experimental apiary (University Paul Sabatier – Toulouse III, France) between January 2019 and August 2020. For the experiments, we used lead (PbCl_2_; CAS #7758-95-4 and PbC_4_H_6_O_4_ 3H2O; CAS #6080-56-4), zinc (ZnCl_2_; CAS # 7646-85-7 and ZnC_4_H_6_O_4_; CAS #557-34-6) and arsenic (NaAsO_2_; CAS #7784-46-5) (all from Sigma-Aldrich, St Louis, MO). The metallic compounds were either dissolved in 30% (w/v) sucrose solution (for feeding, proboscis response and electrophysiological assays) or in mineral water (for antennal response and electrophysiological assays). We tested both chloride or acetate salts of lead (Pb) and zinc (Zn). For arsenic (As) (for the sake of simplicity, we will refer to it as a metal pollutant), we chose arsenite AsIII, a highly toxic form [35] derived from smelting and found in insecticides [36,37]. We used nominal concentrations of 0.001, 0.013, 0.129 and 12.83 μM of As; 0.36, 3.60, 35.96 μM and 3.6 mM of Pb; and 0.012, 0.12, 1.22 and 122.3 mM of Zn (see Table S1 for correspondences in ppm and mg.L^-1^). The three lower concentrations of Zn and Pb and all concentrations of As have been reported in field studies of metal pollution (Table S1). The highest concentrations were above the regulatory levels in food as defined by the WHO [38,39], and were assessed through chemical analysis, which gave a good recovery rate (Table S1).

### (b) Feeding assays

We tested the ability of bees to discriminate metal salts in food in assays in which groups of bees could self-select foods over several hours [40]. We collected workers of unknown age at the colony entrance of five different hives, as they returned from foraging. The bees were cold-anaesthetized and placed in groups of 20 in plastic cages (80 × 50 × 40 mm), for 3 days in an incubator (dark, 28 °C ± 1°C, 60% relative humidity). Each cage contained two 2 mL feeding vials (Eppendorf) pierced with two 2 mm holes at the bottom to allow drinking of the sucrose solutions they contained. In the no-choice condition, bees were offered only one type of food: either 30% sucrose solution or 30% sucrose solution containing either As, Pb or Zn salts at one of the concentrations in Table S1. In the choice condition, bees were offered one vial containing pure sucrose and one vial containing a sucrose and metal salt solution. Feeding vials were weighed prior to be placed in the experimental cages, then removed, weighed and replaced by fresh ones every 24 h during 3 days. Cages without bees were used to measure the evaporation rate from the feeding vials. The amount of solution consumed daily was estimated by measuring weight loss in each vial every 24 h. The average value for evaporation of each treatment was subtracted from this final value for each vial. The number of dead bees in each cage was counted every hour (from 9 am to 5 pm), thus allowing the calculation of the mean daily consumption per bee (daily consumption divided by the mean number of bees alive in the cage).

### (c) Devaluation assays following antennal stimulation

We tested the ability of bees to perceive metal salts diluted in water using a devaluation assay that assesses whether repeatedly pairing a previously rewarding odour to contaminated water delivered to the antennae could lead to the devaluation of this odour, thus meaning that it would be perceived as aversive [20]. Workers of unknown age were collected from the top of the frames of eight different hives, cooled on ice, and harnessed in individual plastic holders allowing free movements of their antennae and mouthparts. We fixed their head to the holder using a droplet of melted bee wax, fed them 5 μL of sucrose solution (50% w/v) and let them rest for 3 h in an incubator (dark, 28 ± 1 °C, 60% relative humidity). Before starting the experiment, we checked for intact proboscis extension reflex (PER) by touching the antennae with a toothpick soaked in 50% (w/v) sucrose solution without subsequent feeding. Bees that did not exhibit the reflex were discarded.

The first phase of the assay started with three trials pairing an odour (pure 1-nonanol) with a 50% (w/v) sucrose solution reward. The second phase consisted of 10 trials where a presentation of the trained odorant was followed by the stimulation of the antennae with a metal solution, or just water for the control group. For the second phase of the assay, we only kept bees that performed a PER response to 1-nonanol (92% of the bees). The odour was presented via an automated odour delivery system with a continuous air-stream as described in [41]. For each trial, the harnessed bee was placed in the conditioning set-up for 15 s to allow familiarization, then 1-nonanol was released for 6 s. Four seconds after odour onset, the antennae were stimulated with 50% (w/v) sucrose solution (phase 1), or metal solutions, or water (phase 2) for 2 s followed by 1 s of feeding with sucrose. The bee was left in the conditioning setup for 20 s before being removed. Inter-trial interval was 15 min for both phases. We recorded the proboscis extension response at each trial (extension = 1, no extension = 0).

### (d) Devaluation assay following proboscis stimulations

We assessed whether bees were able to perceive metal salts through their proboscis and/or post-ingestive consequences (i.e. malaise-like state) [23], by testing their potential devaluating effect when applied to the proboscis. We collected workers of unknown age from the top of the frames of four different hives, harnessed and fed them with 5 μL of sucrose solution (50% w/v) and left them to rest for 3 h (dark, 28 ± 1°C, 60% relative humidity). For 12 trials, bees were conditioned to associate 1-nonanol with ingestion of the sucrose-contaminated stimulus: after application of a droplet of 30% (w/v) sucrose onto the antennae to trigger PER, a 0.4 μL droplet of a metal-spiked sucrose solution was delivered to the proboscis. We recorded the proboscis extension response upon odour delivery for each of the 12 trials (extension = 1, no extension = 0). Here again, we expected that any decrease of response frequency would reveal aversion to some stimuli. In addition, we collected bees in the same conditions, harnessed and fed them with 4.8 μL (12 times 0.4 μL) of each solution and monitored their survival for 150 min (i.e. the duration of the proboscis response assay).

### (e) Electrophysiological recordings

We performed electrophysiological recordings on chaetic sensilla [42,43], which can be easily identified by their external morphology [44] (Figure 3A). We focused on the antennae, the organs concentrating the highest number of taste sensilla [29], and specifically on the tip ventral zones [45], which are devoid of olfactory sensilla [42].

We immobilized the antennal flagellum with a metal thread stuck with wax and placed a glass electrode (ext. diameter 10−20 µm) over a single taste sensillum [29]. We used a silver wire inserted into the contralateral eye as grounded reference electrode. Electrodes were pulled from borosilicate glass capillaries, filled with different solutions and stored in a humid chamber before use. We prepared 30 mM sucrose solutions, contaminated or not with metal pollutant salts (Table S1) in 1 mM KCl, which ensured the necessary conductivity for recording, and kept them at 4 °C (1mM KCl was used as the reference). We stimulated taste sensilla in the following order: 1mM KCl, 30 mM sucrose, and 30 mM sucrose containing increasing concentrations of metal pollutant salts. In a separate experiment, increasing concentrations of KCl (1 mM, 10 mM, 50 mM and 500 mM) diluted in 30 mM sucrose, were also tested. All stimuli were applied for 2 s, with an interstimulus interval of 1 min. The recording and reference electrodes were connected to a preamplifier (TasteProbe - Synthec, Kirchzarten, Germany). The electric signals were amplified (×10) using a signal connection interface box (Syntech, Kirchzarten, Germany) in conjunction with a 100-3000Hz band passfilter. Experiments started when the recording electrode contacted the sensillum under study, which triggered data acquisition and storage on a hard disk (sampling rate: 10kHz). We then analysed these data using Autospike (Syntech) and quantified the number of spikes after stimulus onset.

### (f) Statistical analysis

We performed all statistical analyses in R [46]. For the choice assay, we analysed the consumption preference (difference between mean daily consumptions of each food in g/bee) with linear mixed effect models (LMMs; lme4 package [47]), against zero (no preference). For the no-choice assay, we analysed the daily consumption (g/bee) with LMMs. Models were fitted with treatment as a fixed effect and cages nested in hive as a random effect. Models were followed by pairwise comparisons (multcomp package [48]). We analysed the survival probability over three days using a Cox regression model [49].

For antennal and proboscis response assays, we scored the PER of each bee as a binary variable (response=1, no response=0), and analysed the mean score (averaged over the trials) using a binomial generalised linear mixed model (GLMM, lme4 package [47]), with treatment as fixed effect, trial number as a covariate, individual identity nested in the colony, and trial as random grouping variable. For proboscis responses, we also applied GLMMs separately for each trial, with treatment as fixed effect and individual identity nested in the colony, to better capture the temporal dynamics of responses. We analysed the survival probability over 150 min using a Cox regression model.

Electrophysiological data were analysed by comparing frequencies of recorded spikes using a negative binomial GLMM using Template Model Builder [50], with treatment as a fixed effect and bee identity as random variable to take into account the repeated measurements per individual. Models were followed by pairwise comparisons [48].

## 3. Results

### (a) Bees only avoided high concentrations of Pb and Zn in food

The highest concentrations of Zn salts (both chloride and acetate) in food were toxic, inducing high mortality after 24 h (Cox model: p<0.001 and p=0.010 respectively, Figure S1). Therefore, we compared food consumption across all treatments and for choice and no-choice feeding assays over the first 24 h only (Figure S2).

We first tested whether bees discriminated metals in food when given a choice between two accessible sucrose solutions, one of which contained one out of four concentrations of As, Pb or Zn (Figure 1A). None of the As solutions were avoided or preferred when compared to pure sucrose solution. Similarly, there was no difference in consumption of sucrose solutions containing low concentrations of Pb and Zn and pure sucrose (Table S2A). However, the highest concentrations of Pb (3.6 mM, LMM: p<0.001 for both chloride and acetate) and Zn (122.3 mM, LMM: p<0.001 for both salts) were consumed significantly less than pure sucrose.

**Figure 1:**
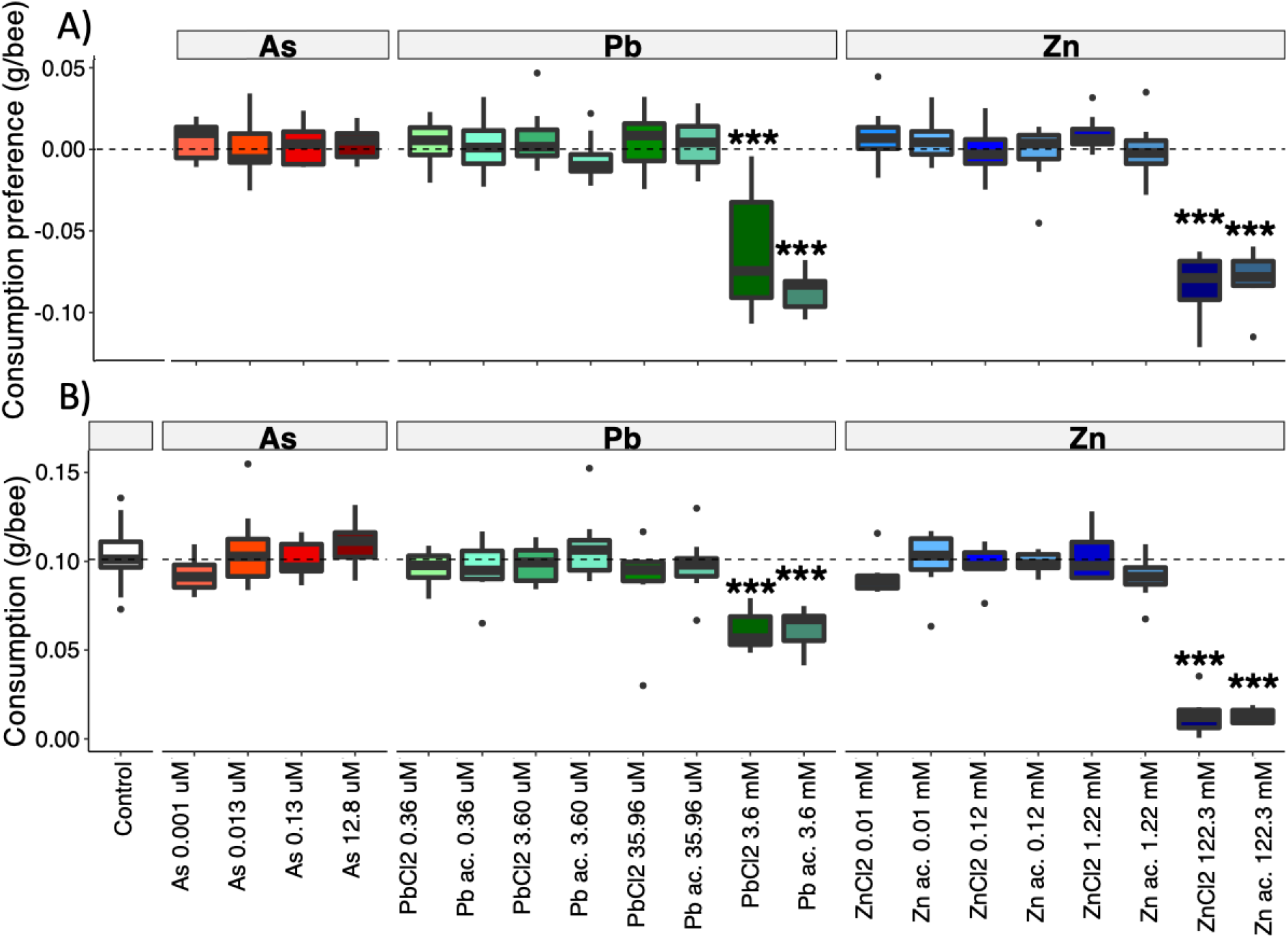
Feeding assays. **A) Choice experiment.**Consumption preferences (difference in daily consumption between the two solutions) are plotted. Positive values: preference for the metal pollutant solution; zero (dotted line): no preference; negative values: preference for the pure sucrose solution. N = 8 cages of 20 bees per treatment. **B) No-choice experiment**. Daily food consumption of each solution; the dotted line indicates the median value for control bees (plain sucrose solution, white). N = 8 cages per treatment and N = 27 cages for control bees. In both experiments we used three metals (arsenic - red, lead - green, zinc - blue) at four concentrations each. Box plots show median (horizontal line), 25th to 75th percentiles (box), smallest and highest values within 1.5*inter-quartile range of the hinge (error bars), and outliers (dots). *p<0.05, **p<0.01, ***p<0.001: differences with zero (A) or control bees (B), LMMs (Table S2).

We then tested whether bees would still avoid their consumption of metals in food when they had no alternative choice (Figure 1B). Bees showed similar consumption of food containing either As (all concentrations), low concentrations of Pb and Zn, or no metal salts (control) (LMM: p>0.05). However, they reduced their total food consumption by 40% when it contained the highest concentration of Pb (3.6 mM, LMM: p<0.001 for both salts), and by 87% when it contained the highest concentration of Zn (122.3 mM, LMM: p<0.001 for both salts) (Table S2B). These effects were independent of the chemical forms (acetate or chloride) of Pb and Zn (LMM: p>0.05 for pairwise comparison for each concentration). For all metals, we found a positive significant correlation between the quantity of metal ingested and the quantity of metal bioaccumulated in the bees’ bodies (As: R= 0.91, p<0.001; Pb: R=0.97, p<0.001; Zn: R=0.82, p<0.001; Figure S3), hence validating our exposure protocol.

### (b) Bees perceived only high concentrations of the three metals, with their antennae and proboscis

We tested whether bees were able to perceive metal salts through their antennae in a devaluation experiment. Since the conditioning odour had been associated with sucrose, we expected a progressive decrease of the rate of conditioned PER over subsequent unrewarded (water) presentations in all groups. If bees perceive metal salts in water, the decrease of response to the metal solution should be stronger than with water. Overall, antennal stimulation with solutions containing metal pollutant salts affected PER responses (Figure 2A, Table S3). The mean PER rate was significantly reduced for the two highest concentrations of As (12.8 μM, 0.13 μM), Zn chloride and acetate (122.3 mM, 1.22 mM), Pb acetate (3.6 mM, 35.96 μM) and only for the second highest concentration of Pb chloride (3.6 mM). We found no overall effect of the chemical form (acetate or chloride) of Pb or Zn (Binomial GLMM: p>0.05 for pairwise comparisons for each concentration). Therefore, bees perceived the highest concentrations of each metal salt through their antennae and reduced their appetitive response.

**Figure 2:**
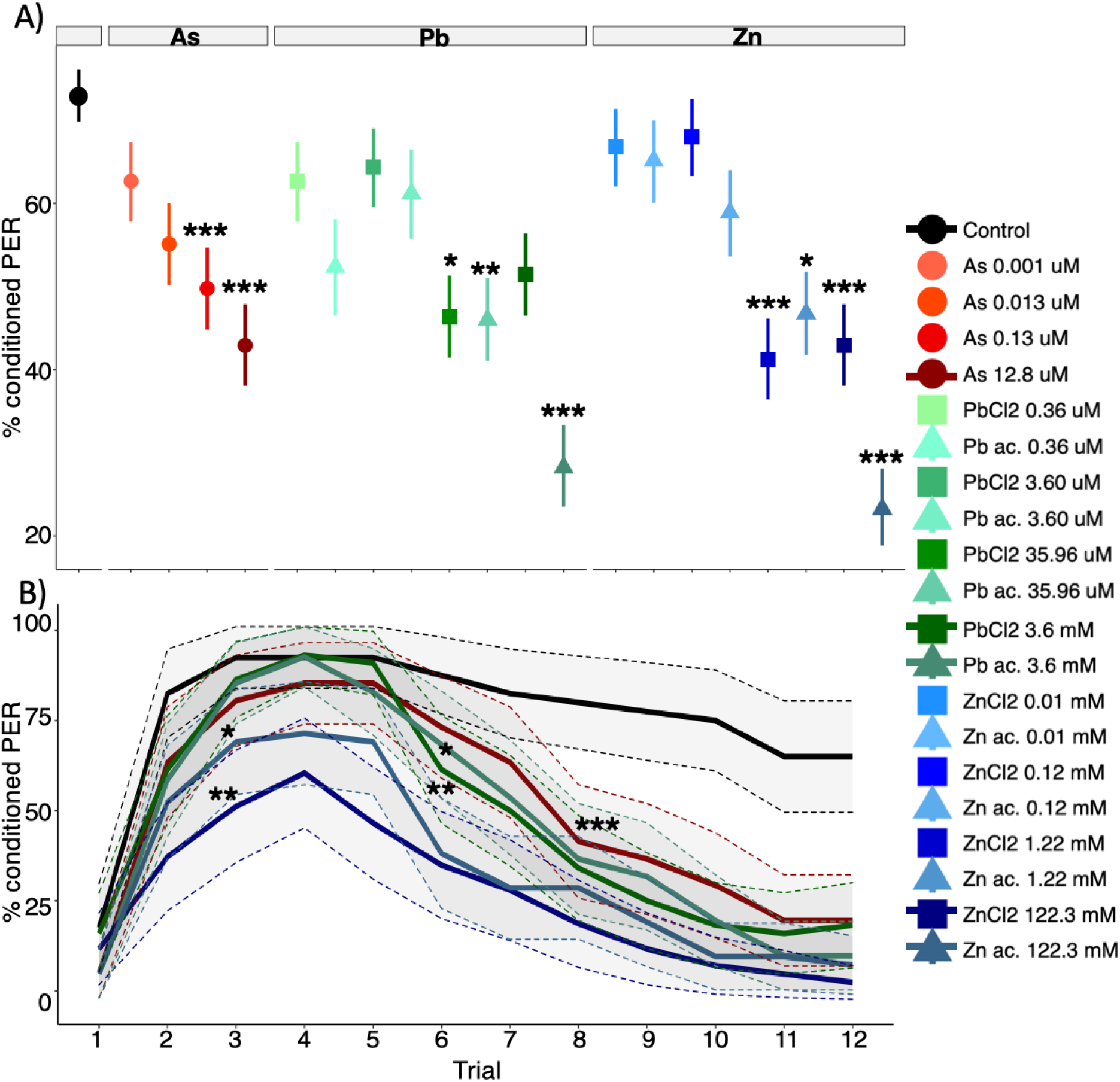
Devaluating effects of metal salts. Mean conditioned proboscis extension response (PER) and 95% confidence intervals (bars in A, shaded in B) across devaluation trials, for each treatment. **A) Application on the antennae**. For lead and zinc, chemical forms are shown by the mean point shape, square for chloride (Cl_2_) and triangle for acetate (C_4_H_6_O_4_). N = 35-41 bees/treatment. *p<0.05, **p<0.01, ***p<0.001: binomial GLMM, compared to controls (N = 79)). **B) Application on the proboscis**. N = 40-42 bees/treatment. *p<0.05, **p<0.01, ***p<0.001: differences with controls (N=40), displayed only for the first trial showing significant differences (binomial GLMM).

**Figure 3:**
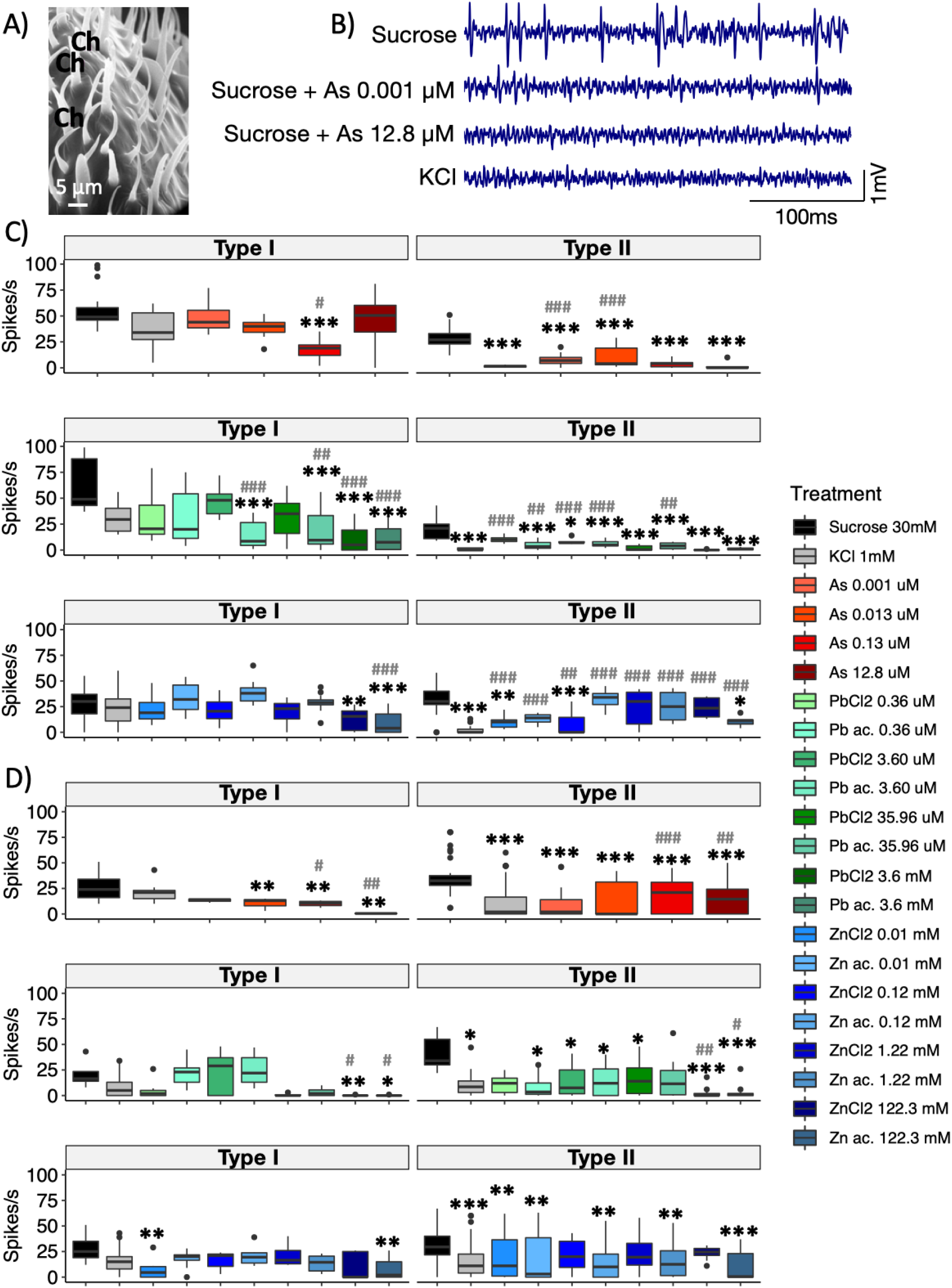
Electrophysiological recordings of the gustatory neurons from the antennae. **A)** Scanning electron microscope picture of the surface of the antenna showing chaetic sensilla (Ch) chosen for recording. **B)** Examples of spike trains recorded from a type II sensilla in response to various stimuli. Note the decreased spike frequency induced by the presence of As in the sucrose solution. **C, D)** Boxplots of the spiking responses to sucrose (black), KCl (grey), and increasing concentrations of arsenic (red), lead (green) and zinc (blue), for a type I sensilla (responding to both KCL and sucrose, left) or a type II (responding to sucrose only, right). **C)** Stimulation with metal salts diluted in sucrose solution (As: N = 4; Pb: N = 5; Zn: N = 6). **D)** Stimulation with metal salts diluted in water (As: N = 4; Pb: N = 5; Zn: N = 6). (#/*p<0.05, **p<0.01, ***p<0.001: differences with sucrose (star) or KCl (hash) (pairwise comparisons following GLMM).

We assessed whether bees were able to perceive metal salts through their proboscis and/or post-ingestive consequences (i.e. malaise-like state) using another devaluation paradigm, in which bees were trained to associate an odour with a sucrose presentation on the antennae (to induce PER) followed by delivery of a lower concentration of sucrose to their proboscis. Metals were diluted in sucrose instead of water to ensure their ingestion. Bees that received sucrose containing metals to the proboscis reduced their PER response more than controls (Figure 2B). Zn-treated bees showed significantly lower levels of PER as early as the 3^rd^ trial (Binomial GLMM: p=0.001 for Zn chloride, p=0.036 for Zn acetate). By contrast, the response levels of Pb and As groups initially reached a maximum similar to the controls. These response levels decreased from the 6^th^ trial onwards with Pb (Binomial GLMM: p=0.009 for chloride, p=0.044 Pb acetate), and from the 8^th^ trial onwards for As (Binomial GLMM: p<0.001). These effects were independent of the chemical forms of Zn and Pb (Tukey HSD: p>0.05). Thus, bees seemed to evaluate negatively all three metals, through their proboscis and/or post-ingestive effects, as eventually responded to all contaminated solutions by markedly decreased PER rates (GLMM: mean PER response: p<0.001 for all treatments). The ingested volumes of metal pollutant solutions were not sufficient to impact their survival over the duration of the experiment (Figure S4).

### (c) Highly concentrated metals inhibit sucrose-evoked activity in taste receptors

We finally performed electrophysiological recordings to investigate the mechanisms by which bees detect metal salts, focusing on neurons in antennal gustatory sensilla (Figures 3A-B), which are mostly tuned to detect sugars and salts [29]. Irrespective of responses to metal pollutant salts, we identified two main response profiles that lead us to distinguish two functional categories of sensilla: those responding to both sucrose and KCl (Type I, 722 recordings) and those responding to sucrose only (Type II, 953 recordings).

We recorded electrophysiological responses to ascending concentrations of each metal pollutant salt diluted in 30% mM sucrose (Figure 3C). Some sensilla responded equally to both sucrose and KCl (Type I sensilla), but showed a drop in spike frequency in response to high metal concentrations. This is a specific response to metal salts since adding a nutrient salt such as KCl to sucrose had the opposite effect (Figure S5). Other sensilla responded much more to sucrose than to KCl (Type II sensilla) and showed a similar reduction in their activity in response to all metal pollutant salts, when compared to pure sucrose. Overall, the chemical form of Pb or Zn had no effect (GLMM: p>0.05; except for 3.60 μM Pb on type I sensilla: p<0.001; and 0.12m Zn on type II sensilla: p<0.001). Thus, the presence of metal pollutant salts at high levels in sucrose solution could be detected by antennal gustatory neurons, which reduced their activity.

We then asked whether metal salts could be detected independently of the presence of sucrose, and thus used water solutions as stimuli (Figure 3D). Type I sensilla responded to low concentrations of all metal salts similarly to KCl or sucrose. By contrast, they reduced their spike frequency when stimulated with high metal concentrations, as compared to both KCl and sucrose. Type II sensilla failed to show marked activity in response to most metal solutions, as they did for KCl. Thus, metal salts did not trigger a specific response pattern by themselves, but rather reduced sucrose-triggered responses when added at high concentration to the sucrose solution.

## 4. Discussion

Pollinators are impacted by metal pollutants [5,18,51,52]. Here we showed that bees have only a limited capacity to detect and avoid these poisons in food. Honey bees perceived very high (unrealistic) concentrations, of Pb and Zn through their proboscis and antennae, and avoided ingesting them. Sucrose containing concentrated As was detected, but still consumed. Lower, yet harmful, field-realistic concentrations of the metal pollutants were neither avoided nor detected in our experimental conditions. Electrophysiological recordings from gustatory neurons confirmed that bees can only taste a limited concentration range of these metal pollutants.

Bees avoided Zn and Pb (but not As) at high concentrations above most environmental levels, even in the absence of alternative food sources. This observation is consistent with previous reports of decreased food consumption following exposure to high Zn or Pb levels [15,18,53]. However, honey bees ingested sucrose solutions containing all three metal salts at concentrations similar to those found in nectar [54,55]. While Zn at low concentrations is an essential micronutrient, Pb and As can be highly toxic [36]. The absence of any behavioural responses to these solutions at field relevant concentrations thus suggests honey bees are incapable of discriminating between essential and toxic metals. This is consistent with studies reporting indiscriminate visits on metal-contaminated flowers [19,31,56]. At these realistic concentrations, metal pollutants alter the development [15], learning and memory [51,52], the metabolism [16,17] and antioxidative responses [57].

Stimulations of gustatory organs with metal solutions showed high concentrations were perceived through the antennae and the proboscis. This devaluating effect occurred with antennal stimulation only. It is thus independent of post-ingestive effects such as those observed with other toxic substances [20,23,24]. While methodological differences make difficult a direct comparison of devaluation responses with and without ingestion, they did not seem to be much stronger when metals were delivered to the proboscis and ingested, indicating that post-ingestion effects (if any) would have been minimal.

The detection of metals by taste receptors was sufficient to reduce appetitive behaviour. The decreased responsiveness to repeated stimulations with contaminated sucrose on taste receptors likely results from a mismatch between expected and obtained rewards, possibly because peripheral detection of metals actively inhibited appetitive behaviour and/or because sucrose-sensitive taste receptors were inhibited. Both mechanisms have been involved in the feeding suppression triggered by plant-derived deterrents [26], but electrophysiological data was lacking to confirm the implication of either process in these and previous behavioural effects of metal pollutants [18,53]. Here, concentrated Pb, As and Zn decreased sucrose-evoked spike frequencies in bee taste receptors’ response to sucrose. Importantly, this effect was due to the metals themselves as it was observed irrespective of the salts used (acetate or chloride), and in a different concentration range as for common salts (e.g. KCl). By contrast, we found no clear evidence of specific detection systems, consistently with the limited molecular repertoire of gustatory receptors in this species [58]. Thus, reduced neural activity might result from non-specific toxic effects, such as oxidative stress and ion channel dysfunction [59,60]. While the exact mechanism remains to be determined, very high metal concentrations rarely encountered (even in polluted environments) can trigger rejection of feeding sites that would be toxic at short term, as already observed for naturally deterrent compounds (e.g. bitter substances) [21,61]. However, since it is not observed for field–relevant doses of metals, it is unlikely to have been selected, contrary to the anti-feeding action of many phytochemicals acting as a plant defence mechanism against phytophagous insects [26].

## 4. Conclusion

Our study echoes to the recent findings that bees cannot detect harmful insecticides through taste [40,62]. It calls for further research to better characterize the response of bees to heavy metal pollutants, including to combinations of different elements [56]. Since metal pollutants are undetected and consumed by bees, low amounts can bioaccumulate, which may lead to long-term detrimental effects on individuals and colony health [63]. Evidence of hazards of heavy metals on terrestrial wildlife worryingly accumulate [2]. It has become an urgent issue to account for such effects in order to adjust permissible levels of environmental metal pollution accordingly [2].

## Supporting information

Supplementary ables 1-3 an Figures 1-5-

Raw data

## Acknowledgements

We thank Olivier Fernandez for beekeeping and Cristian Pasquaretta for help with the statistical analyses. We thank Camille Duquenoy (Service ICP-MS Observatoire Midi-Pyrénées) for her assistance in the lab.

## Author contribution

CM, MGBS, AE, ABB, ML and JMD designed the study. CM, MGBS, LL, OB, GM, JS, GLR and DB collected the data. CM analysed the data and wrote the first draft of the manuscript. All authors reviewed the manuscript.

## Data availability

Raw data are available in Dataset S1.

## References

1. Potts SG, Biesmeijer JC, Kremen C, Neumann P, Schweiger O, Kunin WE. 2010 Global pollinator declines: trends, impacts and drivers. Trends Ecol. Evol. 25, 345–353. (doi:10.1016/j.tree.2010.01.007)

2. Monchanin C, Devaud J-M, Barron A, Lihoreau M. 2021 Current permissible levels of heavy metal pollutants harm terrestrial invertebrates. Sci. Total Environ. 779, 146398. (doi:10.1016/j.scitotenv.2021.146398)

3. Bradl HB. 2005 Sources and origins of heavy metals. In Interface Science and Technology (ed HB Bradl), pp. 1–27. Elsevier. (doi:10.1016/S1573-4285(05)80020-1)

4. Nriagu JO, Pacyna JM. 1988 Quantitative assessment of worldwide contamination of air, water and soils by trace metals. Nature 333, 134–139.

5. Thimmegowda GG, Mullen S, Sottilare K, Sharma A, Mohanta SS, Brockmann A, Dhandapany PS, Olsson SB. 2020 A field-based quantitative analysis of sublethal effects of air pollution on pollinators. Proc. Natl. Acad. Sci. USA 117, 20653–20661. (doi:10.1073/pnas.2009074117)

6. Gutiérrez M, Molero R, Gaju M, van der Steen J, Porrini C, Ruiz JA. 2020 Assessing heavy metal pollution by biomonitoring honeybee nectar in Córdoba (Spain). Environ. Sci. Pollut. Res. 27, 10436–10448. (doi:10.1007/s11356-019-07485-w)

7. Roman A. 2007 Content of some trace elements in fresh honeybee pollen. Pol. J. Food Nutr. 57, 475–478.

8. Balestra V, Celli G, Porrini C. 1992 Bees, honey, larvae and pollen in biomonitoring of atmospheric pollution. Aerobiologia 8, 122–126. (doi:10.1007/BF02291339)

9. Goretti E et al. 2020 Heavy metal bioaccumulation in honey bee matrix, an indicator to assess the contamination level in terrestrial environments. Environ. Pollut. 256, 113388. (doi:10.1016/j.envpol.2019.113388)

10. Satta A, Verdinelli M, Ruiu L, Buffa F, Salis S, Sassu A, Floris I. 2012 Combination of beehive matrices analysis and ant biodiversity to study heavy metal pollution impact in a post-mining area (Sardinia, Italy). Environ. Sci. Pollut. Res. 19, 3977–3988. (doi:10.1007/s11356-012-0921-1)

11. Zhou X, Taylor MP, Davies PJ, Prasad S. 2018 Identifying sources of environmental contamination in European honey bees (Apis mellifera) using trace elements and lead isotopic compositions. Environ. Sci. Technol. 52, 991–1001. (doi:10.1021/acs.est.7b04084)

12. Domingo JL. 1994 Metal-induced developmental toxicity in mammals: a review. J. Toxicol. Environ. Health 42, 123–141. (doi:10.1080/15287399409531868)

13. Saulnier A, Bleu J, Boos A, El Masoudi I, Ronot P, Zahn S, Del Nero M, Massemin S. 2020 Consequences of trace metal cocktail exposure in zebra finch (Taeniopygia guttata) and effect of calcium supplementation. Ecotoxicol. Environ. Saf. 193, 110357. (doi:10.1016/j.ecoenv.2020.110357)

14. Tchounwou PB, Yedjou CG, Patlolla AK, Sutton DJ. 2012 Heavy metal toxicity and the environment. In Molecular, Clinical and Environmental Toxicology (ed A Luch), pp. 133–164. Basel: Springer Basel. (doi:10.1007/978-3-7643-8340-4_6)

15. Di N, Hladun KR, Zhang K, Liu T-X, Trumble JT. 2016 Laboratory bioassays on the impact of cadmium, copper and lead on the development and survival of honeybee (Apis mellifera L.) larvae and foragers. Chemosphere 152, 530–538. (doi:10.1016/j.chemosphere.2016.03.033)

16. Nikolić TV, Kojić D, Orčić S, Batinić D, Vukašinović E, Blagojević DP, Purać J. 2016 The impact of sublethal concentrations of Cu, Pb and Cd on honey bee redox status, superoxide dismutase and catalase in laboratory conditions. Chemosphere 164, 98–105. (doi:10.1016/j.chemosphere.2016.08.077)

17. Nikolić TV, Kojić D, Orčić S, Vukašinović EL, Blagojević DP, Purać J. 2019 Laboratory bioassays on the response of honey bee (Apis mellifera L.) glutathione S-transferase and acetylcholinesterase to the oral exposure to copper, cadmium, and lead. Environ. Sci. Pollut. Res. 26, 6890–6897. (doi:10.1007/s11356-018-3950-6)

18. Burden CM, Morgan MO, Hladun KR, Amdam GV, Trumble JJ, Smith BH. 2019 Acute sublethal exposure to toxic heavy metals alters honey bee (Apis mellifera) feeding behavior. Sci. Rep. 9, 4253. (doi:10.1038/s41598-019-40396-x)

19. Chicas-Mosier AM, Cooper BA, Melendez AM, Pérez M, Oskay D, Abramson CI. 2017 The effects of ingested aqueous aluminum on floral fidelity and foraging strategy in honey bees (Apis mellifera). Ecotoxicol. Environ. Saf. 143, 80–86. (doi:10.1016/j.ecoenv.2017.05.008)

20. Ayestaran A, Giurfa M, de Brito Sanchez MG. 2010 Toxic but drank: gustatory aversive compounds induce post-ingestional malaise in harnessed honeybees. PLoS ONE 5, e15000. (doi:10.1371/journal.pone.0015000)

21. de Brito Sanchez MG, Giurfa M, de Paula Mota TR, Gauthier M. 2005 Electrophysiological and behavioural characterization of gustatory responses to antennal ‘bitter’ taste in honeybees. Eur. J. Neurosci. 22, 3161–3170. (doi:10.1111/j.1460-9568.2005.04516.x)

22. Guiraud M, Hotier L, Giurfa M, de Brito Sanchez MG. 2018 Aversive gustatory learning and perception in honey bees. Sci. Rep. 8, 1343. (doi:10.1038/s41598-018-19715-1)

23. Wright GA, Mustard JA, Simcock NK, Ross-Taylor AAR, McNicholas LD, Popescu A, Marion-Poll F. 2010 Parallel reinforcement pathways for conditioned food aversions in the honeybee. Curr. Biol. 20, 2234–2240. (doi:10.1016/j.cub.2010.11.040)

24. Lai Y, Despouy E, Sandoz J-C, Su S, de Brito Sanchez MG, Giurfa M. 2020 Degradation of an appetitive olfactory memory via devaluation of sugar reward is mediated by 5-HT signaling in the honey bee. Neurobiol. Learn. Mem. 173, 107278. (doi:10.1016/j.nlm.2020.107278)

25. Herbert EW, Shimanuki H. 1978 Mineral requirements for brood-rearing by honeybees fed a synthetic diet. J. Apic. Res. 17, 118–122. (doi:https://doi.org/10.1080/00218839.1978.11099916)

26. Koul O. 2008 Phytochemicals and insect control: an antifeedant approach. Crit. Rev. Plant Sci. 26, 1–24. (doi:10.1080/07352680802053908)

27. Hodgson ES. 1957 Electrophysiological studies of arthropod chemoreception. Responses of labellar chemoreceptors of the blowfly to stimulation by carbohydrates. J. Insect Physiol. 1, 240–247. (doi:10.1016/0022-1910(57)90039-2)

28. Schoonhoven LM, Jermy T. 1977 A behavioural and electrophysiological analysis of insect feeding deterrents. In Crop protection agents-their biological evaluation (ed NR McFarlane), pp. 133–147. New York, NY: Academic Press.

29. de Brito Sanchez MG. 2011 Taste perception in honey bees. Chem. Senses 36, 675–692. (doi:10.1093/chemse/bjr040)

30. Hladun KR, Smith BH, Mustard JA, Morton RR, Trumble JT. 2012 Selenium toxicity to honey bee (Apis mellifera L.) pollinators: effects on behaviors and survival. PLoS ONE 7, e34137. (doi:10.1371/journal.pone.0034137)

31. Sivakoff FS, Gardiner MM. 2017 Soil lead contamination decreases bee visit duration at sunflowers. Urban Ecosyst. 20, 1221–1228. (doi:10.1007/s11252-017-0674-1)

32. Xun E, Zhang Y, Zhao J, Guo J. 2018 Heavy metals in nectar modify behaviors of pollinators and nectar robbers: consequences for plant fitness. Environ. Pollut. 242, 1166–1175. (doi:10.1016/j.envpol.2018.07.128)

33. Brodschneider R, Crailsheim K. 2010 Nutrition and health in honey bees. Apidologie 41, 278–294. (doi:10.1051/apido/2010012)

34. ATSDR. 2019 The ATSDR 2019 Substance Priority List. See https://www.atsdr.cdc.gov/spl/index.html (accessed on 13 May 2020).

35. Tyson J. 2013 The determination of arsenic compounds: a critical review. ISRN Anal. Chem. 2013, 1–24. (doi:10.1155/2013/835371)

36. Bissen M, Frimmel FH. 2003 Arsenic — a Review. Part I: Occurrence, Toxicity, Speciation, Mobility. Acta Hydrochim. Hydrobiol. 31, 9–18. (doi:10.1002/aheh.200390025)

37. Defarge N, Spiroux de Vendômois J, Séralini GE. 2018 Toxicity of formulants and heavy metals in glyphosate-based herbicides and other pesticides. Toxicol. Rep. 5, 156–163. (doi:10.1016/j.toxrep.2017.12.025)

38. Codex Alimentarius. 1984 Contaminants, Joint FAO/WHO Food standards Program (Vol. XVII, 1st ed.), 163–170.

39. Codex Alimentarius. 2015 Codex general standard for contaminants and toxins in food and feed - CODEX STAN 193-1995.

40. Kessler SC, Tiedeken EJ, Simcock KL, Derveau S, Mitchell J, Softley S, Radcliffe A, Stout JC, Wright GA. 2015 Bees prefer foods containing neonicotinoid pesticides. Nature 521, 74–76. (doi:10.1038/nature14414)

41. Aguiar JMRBV, Roselino AC, Sazima M, Giurfa M. 2018 Can honey bees discriminate between floral-fragrance isomers? J. Exp. Biol. 221, jeb180844. (doi:10.1242/jeb.180844)

42. Esslen J, Kaissling K-E. 1976 Zahl und Verteilung antennaler Sensillen bei der Honigbiene (Apis mellifera L.). Zoomorphologie 83, 227–251. (doi:10.1007/BF00993511)

43. Hodgson ES, Lettvin JY, Roeder KD. 1955 Physiology of a primary chemoreceptor unit. Science 122, 417–418. (doi:10.1126/science.122.3166.418)

44. Whitehead W, Larsen JR. 1976 Electrophysiological responses of galeal contact chemoreceptors of Apis mellifera to selected sugars and electrolytes. J. Insect Physiol. 22, 1609–1616.

45. Haupt SS. 2004 Antennal sucrose perception in the honey bee (Apis mellifera L.): behaviour and electrophysiology. J. Comp. Physiol. A 190, 735–745. (doi:10.1007/s00359-004-0532-5)

46. RStudio Team. 2015 RStudio: integrated development for R. RStudio, Inc., Boston, MA URL http://www.rstudio.com/.

47. Bates D, Mächler M, Bolker B, Walker S. 2015 Fitting linear mixed-effects models using lme4. J. Stat. Softw. 67, 1–48. (doi:10.18637/jss.v067.i01)

48. Hothorn T, Bretz F, Westfall P. 2008 Simultaneous inference in general parametric models. Biom. J. 50, 346–363.

49. Therneau TM. 2020 coxme: Mixed Effects Cox Models. R package version 2.2-16. https://CRAN.R-project.org/package=coxme.

50. Brooks M E, Kristensen K, van Benthem KJ, Magnusson A, Berg C W, Nielsen A, Skaug H J, Maechler M, Bolker B M. 2017 glmmTMB balances speed and flexibility among packages for zero-inflated generalized linear mixed modeling. R J. 9, 378–400.

51. Monchanin C et al. 2021 Chronic exposure to trace lead impairs honey bee learning. Ecotoxicol. Environ. Saf. 212, 112008. (doi:https://doi.org/10.1016/j.ecoenv.2021.112008)

52. Monchanin C, Drujont E, Devaud J-M, Lihoreau M, Barron AB. 2021 Heavy metal pollutants have additive negative effects on honey bee cognition. J. Exp. Biol. (doi:10.1101/2020.12.11.421305)

53. Teixeira De Sousa R. 2019 Behavioural regulation of mineral salt intake in the adult worker honey bee, Apis mellifera. PhD thesis, Newcastle University, Newcastle, UK. See http://theses.ncl.ac.uk/jspui/handle/10443/4480.

54. Hajar EWI, Sulaiman AZB, Sakinah AMM. 2014 Assessment of heavy metals tolerance in leaves, stems and flowers of Stevia rebaudiana plant. Procedia Environ. Sci. 20, 386–393. (doi:10.1016/j.proenv.2014.03.049)

55. Maiyo WK, Kituyi JL, Mitei YJ, Kagwanja SM. 2014 Heavy metal contamination in raw honey, soil and flower samples obtained from Baringo and Keiyo Counties, Kenya. Int. J. Emerg. Sci. Eng. 2, 5–9.

56. Hladun KR. 2013 Effects of selenium accumulation on phytotoxicity, herbivory, and pollination ecology in radish (Raphanus sativus L.). Environ. Pollut. 172, 70–75. (doi:https://doi.org/10.1016/j.envpol.2012.08.009)

57. Gauthier M, Aras P, Jumarie C, Boily M. 2016 Low dietary levels of Al, Pb and Cd may affect the non-enzymatic antioxidant capacity in caged honey bees (Apis mellifera). Chemosphere 144, 848–854. (doi:10.1016/j.chemosphere.2015.09.057)

58. Robertson HM, Wanner KW. 2006 The chemoreceptor superfamily in the honey bee, Apis mellifera: expansion of the odorant, but not gustatory, receptor family. Genome Res. 16, 1395–1403. (doi:10.1101/gr.5057506)

59. Garza-Lombó C, Pappa A, Panayiotidis MI, Gonsebatt ME, Franco R. 2019 Arsenic-induced neurotoxicity: a mechanistic appraisal. JBIC J. Biol. Inorg. Chem. 24, 1305–1316. (doi:10.1007/s00775-019-01740-8)

60. Marger L, Schubert CR, Bertrand D. 2014 Zinc: an underappreciated modulatory factor of brain function. Biochem. Pharmacol. 91, 426–435. (doi:10.1016/j.bcp.2014.08.002)

61. de Brito Sanchez MG, Lorenzo E, Su S, Liu F, Zhan Y, Giurfa M. 2014 The tarsal taste of honey bees: behavioral and electrophysiological analyses. Front. Behav. Neurosci. 8, 1–16. (doi:10.3389/fnbeh.2014.00025)

62. Arce AN, Ramos Rodrigues A, Colgan TJ, Wurm Y, Gill RJ. 2018 Foraging bumblebees acquire a preference for neonicotinoid-treated food with prolonged exposure. Proc. R. Soc. B Biol. Sci. 285, 20180655. (doi:10.1098/rspb.2018.0655)

63. Klein S, Cabirol A, Devaud J-M, Barron AB, Lihoreau M. 2017 Why bees are so vulnerable to environmental stressors. Trends Ecol. Evol. 32, 268–278. (doi:10.1016/j.tree.2016.12.009)

