## Supplementary ables 1-3 an Figures 1-5- for "Honey bees cannot sense harmful concentrations of metal pollutants in food"

**Supplementary materials**

**Table S1: Comparison of the concentrations tested and field measures.** Theoretical concentrations are given in molarity, ppm and mg.L$^{-1}$. The highest concentrations of metals in sucrose solutions, used for subsequent dilutions, were analysed [1]. Solutions were acidified at 3% $HNO_3$ with ultra-pure 69% $HNO_3$ to avoid precipitation or adsorption in containers and then diluted with a $HNO_3$ 3% solution to reduce the spectral interference and viscosity effects. Solutions were then analysed using Inductively Coupled Plasma Emission Spectroscopy (ICP-OES, quantification limit: 5-20 µg.kg$^{-1}$, precision measure: 1-5%; AMETEK Spectro ARCOS FHX22, Kleve, Germany). Mean (minimal-maximal) concentrations of arsenic, lead and zinc recorded in honey and flower samples worldwide. ND: not detected. Values in bold show concentrations above the international permissible values in food as per WHO and FAO (As: 0.2 ppm; Pb: 3 ppm; Zn: 60 ppm [2,3])

| Metal | Nominal concentration (molarity) | Actual concentration (molarity) (recovery percentage given) | Nominal concentration (ppm) | Nominal concentration (mg.L$^{-1}$) | Concentration recorded in honey samples (ppm) | Concentration recorded in flower samples (ppm) |
|---|---|---|---|---|---|---|
| As | 0.001 µM | | 0.0001 | 0.000096 | | |
| | 0.013 µM | | 0.001 | 0.00096 | 0.007 (0.003-0.02) [4] | |
| | 0.129 µM | | 0.010 | 0.0096 | 0.015 (0.002-0.03) [5] | 0.098 (0.075-0.12) [6] |
| | 12.83 µM | 8.72 µM (68%) | **0.853** | **0.96** | 0.56 (0.019-1.39) [7]<br><br>0.52 (ND-1.93) [8] | 0.31 [9] |
| Pb | 0.36 µM | | 0.07 | 0.075 | 0.07 (0.01-0.84) [10]<br><br>0.08 (0.03-0.24) [11] | |
| | 3.60 µM | | **0.66** | **0.75** | 0.62 (0.61-0.63) [12] | 0.61 [13] |
| | 35.96 µM | | **6.61** | **7.45** | 0.720 (ND-4.78) [14]<br><br>14.59 (10-18) [15] | 8.05 [16]<br><br>1.53 (0.13-7.68) [15] |

| | | | | | | |
|---|---|---|---|---|---|---|
| | 3.6 mM | $PbCl_2$ 3.83 mM (94%)<br><br>$PbC_4H_6O_4$ 3.06 mM (85%) | **661** | **745** | | |
| | 0.012 mM | | 0.71 | 0.80 | 0.75 (0.04-5.96) [17]<br><br>0.75 (ND-1.43) [18] | 0.42 (0.05-0.63) [13] |
| Zn | 0.12 mM | | 7.09 | 8.00 | 6.39 (1.37-22.15) [14]<br>7.76 (4.17-22.30) [19] | 17.8 (1.15-49.12) [20] |
| | 1.22 mM | | **70.94** | **79.95** | 9.33 (0.23-73.60) [21]<br>43.88 (4.7-174) [22] | 79.0 [23] |
| | 122.3 mM | $ZnCl_2$ 114.4 mM (94%)<br><br>$ZnC_4H_6O_4$ 386.6 mM (71%) | **7094** | **7995** | | |

13

**Table S2: Parameter estimates from the LMMs for the feeding assay after 24h. A)** For the consumption preference (g/bee) of the choice experiment, compared to 0 (i.e. no preference). **B)** For the food consumption (g/bee) of the no-choice experiment compared to control bees. Significant p-values are shown in bold. SE: standard errors.

| | **A) Choice experiment** | | **B) No-choice experiment** | |
|---|---|---|---|---|
| | Estimate ± SE | p-value | Estimate ± SE | p-value |
| As 0.001 µM | 0.0055 ± 0.0064 | 0.393 | -0.0106 ± 0.0053 | 0.918 |
| As 0.013 µM | 0.0006 ± 0.0065 | 0.987 | 0.0038 ± 0.0053 | 1 |
| As 0.13 µM | 0.0026 ± 0.0064 | 0.691 | -0.0034 ± 0.0054 | 1 |
| As 1.8 µM | 0.0040 ± 0.0064 | 0.537 | 0.0061 ± 0.0053 | 0.999 |
| $PbCl_2$ 0.36 µM | 0.0034 ± 0.0064 | 0.598 | -0.0068 ± 0.0053 | 0.999 |
| $PbC_4H_6O_4$ 0.36 µM | 0.0024 ± 0.0064 | 0.707 | -0.0093 ± 0.0054 | 0.981 |
| $PbCl_2$ 3.60 µM | 0.0062 ± 0.0065 | 0.344 | -0.0034 ± 0.0054 | 1 |
| $PbC_4H_6O_4$ 3.60 µM | -0.0057 ± 0.0064 | 0.380 | 0.0006 ± 0.0055 | 1 |
| $PbCl_2$ 35.96 µM | 0.0038 ± 0.0065 | 0.552 | -0.0171 ± 0.0053 | 0.151 |
| $PbC_4H_6O_4$ 35.96 µM | 0.0030 ± 0.0064 | 0.647 | -0.0079 ± 0.0053 | 0.997 |
| $PbCl_2$ 3.6 mM | -0.0629 ± 0.0065 | **<0.001** | -0.0417 ± 0.0053 | **<0.01** |
| $PbC_4H_6O_4$ 3.6 mM | -0.0866 ± 0.0065 | **<0.001** | -0.0428 ± 0.0054 | **<0.01** |
| $ZnCl_2$ 0.01mM | 0.0079 ± 0.0064 | 0.220 | -0.0106 ± 0.0053 | 0.914 |
| $ZnC_4H_6O_4$ 0.01 mM | 0.0051 ± 0.0065 | 0.435 | -0.0057 ± 0.0055 | 1 |
| $ZnCl_2$ 0.12mM | -0.0018 ± 0.0065 | 0.787 | -0.0049 ± 0.0053 | 1 |
| $ZnC_4H_6O_4$ 0.12 mM | -0.0029 ± 0.0064 | 0.655 | -0.0052 ± 0.0053 | 1 |
| $ZnCl_2$ 1.22mM | 0.0093 ± 0.0064 | 0.153 | -0.0041 ± 0.0053 | 1 |
| $ZnC_4H_6O_4$ 1.22 mM | -0.0005 ± 0.0064 | 0.939 | -0.0138 ± 0.0054 | 0.548 |
| $ZnCl_2$ 122.3mM | -0.0839 ± 0.0065 | **<0.001** | -0.0878 ± 0.0053 | **<0.01** |
| $ZnC_4H_6O_4$ 122.3 mM | -0.0791 ± 0.0065 | **<0.001** | -0.0920 ± 0.005 | **<0.01** |

**Table S3: Parameter estimates from the GLMM for the mean proboscis extension response, compared to control bees, of the antennal response assay.** Significant p-values are shown in bold. SE: standard errors.

| | Estimate ± SE | p-value |
|---|---|---|
| As 0.001 µM | -1.6609 ± 0.6752 | 0.631 |
| As 0.013 µM | -2.1685 ± 0.6541 | 0.106 |
| As 0.13 µM | -2.5849 ± 0.6306 | **<0.001** |
| As 1.8 µM | -3.1880 ± 0.6266 | **<0.001** |
| $PbCl_2$ 0.36 µM | -1.5803 ± 0.6734 | 0.717 |
| $PbC_4H_6O_4$ 0.36 µM | -1.7026 ± 0.7482 | 0.766 |
| $PbCl_2$ 3.60 µM | -1.2555 ± 0.6897 | 0.964 |
| $PbC_4H_6O_4$ 3.60 µM | -1.1799 ± 0.7439 | 0.992 |
| $PbCl_2$ 35.96 µM | -2.6603 ± 0.6261 | **<0.001** |
| $PbC_4H_6O_4$ 35.96 µM | -2.4830 ± 0.6751 | **0.034** |
| $PbCl_2$ 3.6 mM | -1.9016 ± 0.6507 | 0.287 |
| $PbC_4H_6O_4$ 3.6 mM | -3.9365 ± 0.6832 | **<0.001** |
| $ZnCl_2$ 0.01mM | -1.3833 ± 0.6765 | 0.893 |
| $ZnC_4H_6O_4$ 0.01 mM | -0.7728 ± 0.7460 | 1 |
| $ZnCl_2$ 0.12mM | -0.9943 ± 0.6923 | 0.998 |
| $ZnC_4H_6O_4$ 0.12 mM | -0.9144 ± 0.7362 | 0.999 |
| $ZnCl_2$ 1.22mM | -3.1825 ± 0.6204 | **<0.001** |
| $ZnC_4H_6O_4$ 1.22 mM | -2.5721 ± 0.6806 | **0.023** |
| $ZnCl_2$ 122.3mM | -3.1551 ± 0.6315 | **<0.001** |
| $ZnC_4H_6O_4$ 122.3 mM | -4.5625 ± 0.6839 | **<0.001** |

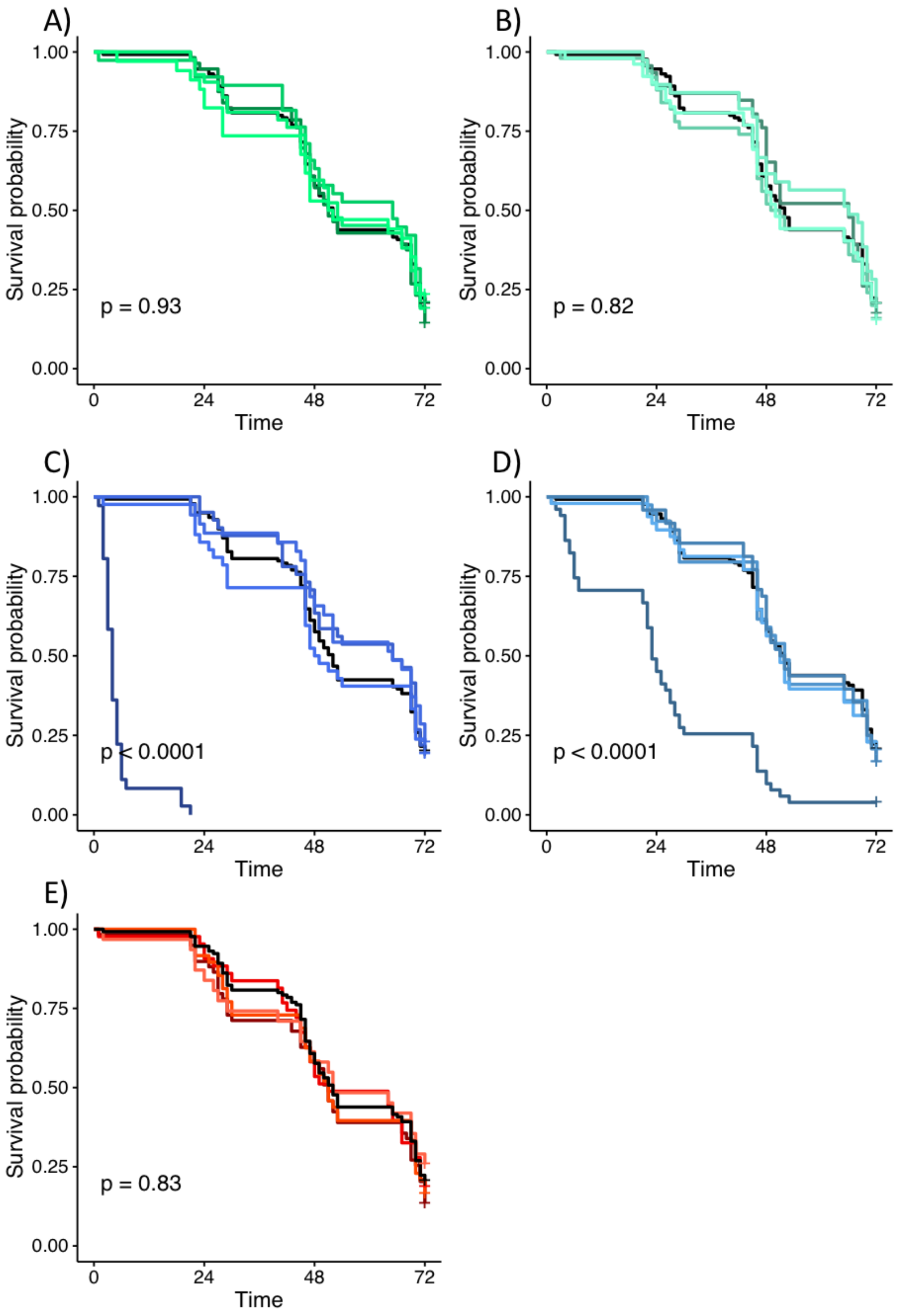


**Figure S1: Survival probability over the 3 days of the no-choice experiment. A)** Lead chloride (0.36 µM-3.6 mM of Pb). **B)** Lead acetate (0.36 µM-3.6 mM of Pb). **C)** Zinc chloride (0.012-122.3 mM of Zn). **D)** Zinc acetate (0.012-122.3 mM of Zn). **E)** Arsenic (0.001-12.83 µM of As). Controls are displayed in black. P-values were obtained from Cox regression models compared to control.

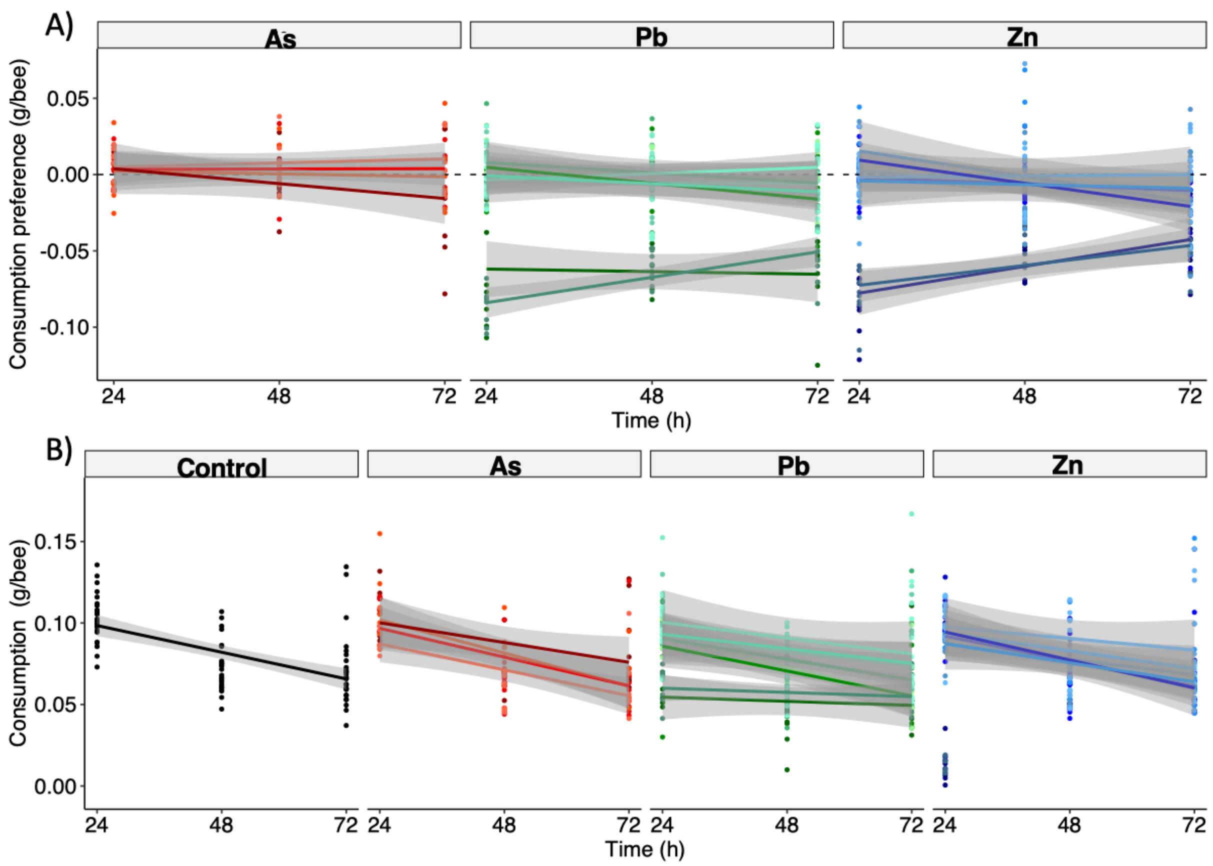


**Figure S2: Feeding assay. A) Choice experiment**. Food consumption preference (g/bee) over the 3 days of experiment. Values over 0 show preference for sucrose-metal diets; values below zero indicate preference for uncontaminated sucrose solution. Dotted line represents no preference. N = 8 cages of 20 bees per treatment **B) No-choice experiment.** Food consumption (g/bee) over the 3 days of experiment. N = 8 cages per treatment and N = 27 cages for control bees. We used three metals (arsenic - red, lead - green, zinc - blue) at four concentrations each.

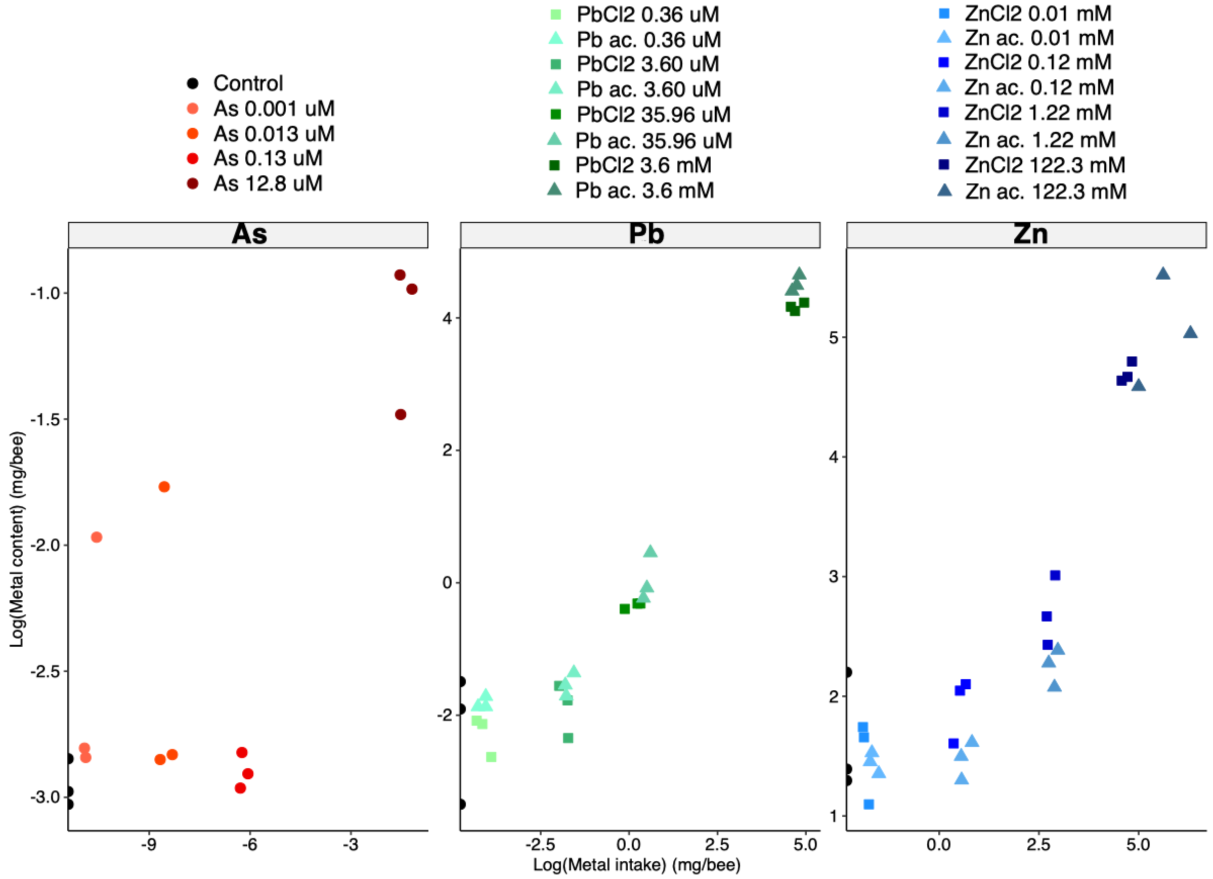


**Figure S3: Metal intake and bioaccumulation in the bodies of bees submitted to the no-choice experiment (log scale).** We used three metals (arsenic - red, lead - green, zinc - blue) in sucrose water at four concentrations each. For lead and zinc, chemical forms are shown by the point shape, square for chloride ($Cl_2$) and triangle for acetate ($C_4H_6O_4$). From the cumulative food consumption over 3 days, and given the metal concentration, we calculated the metal intake per bee (mg/bee). Right after the end of the experiment (3 days), five frozen honey bees were pooled per treatment conditions (N=3 replicates per treatment) and fresh weight was measured. The acid digestion was carried out by adding 2 mL of ultrapure nitric acid (69% w/w; CAS#7697-37-2; optima grade, ThermoFisher Scientific) to the glass vessel containing the bees for 15 min. Then 1 mL of hydrogen peroxide (30% w/w) was added. Glass vessels were introduced in TFM vessels containing 1.5 mL of hydrogen peroxide (35% w/w; CAS# 7722-84-1, Chem Lab), and the mixture was warmed up to 190 °C in a microwave digestion system for 15 min (‘Animal tissue – glass’ settings; MARS 2 Microwave Digestion System, CEM Corporation, USA). The mixture was then cooled and ultrapure water was added to reach a volume of about 25 mL, and weighed. Blank solutions were prepared following the same protocol. Concentrations of metals in honey bee samples were measured using Inductively Coupled Plasma - Mass Spectrometry at Observatoire Midi-Pyrenees ICP-MS platform on a

Thermo ICAP T-Q-ICP-MS (Bremen Germany) (ICP-MS, quantification limit: <0.01µg.kg$^{-1}$, precision measure: 5%). The accuracy of the analytical method was controlled using certified reference materials: lobster hepatopancreas TORT-2, dogfish liver DOLT-3 (LGC Standards, Molsheim, France), caprine horn NYS-RM (New York State Department of Health, USA).

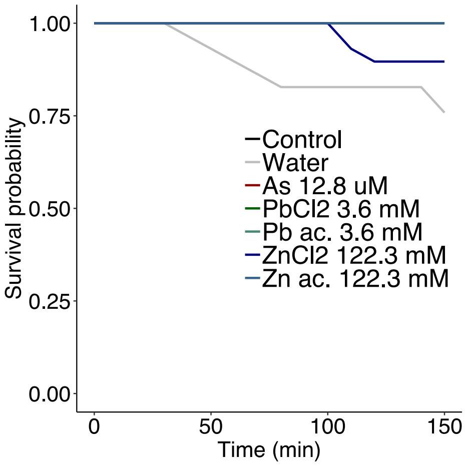


**Figure S4: Survival probability over the duration of the proboscis response assay.** Bees were fed 4.8 µL (equivalent of 0.4 µL ingested during each of the 12 trials) of solutions. As, Pb and Zn acetate treatments had no effect on survival. Bees exposed to Zn chloride exhibited mortality, but not different from the control bees. Bees fed with water only exhibited the highest mortality rate (Cox regression models: $p<0.05$). Note that some of the 7 curves are superposed.

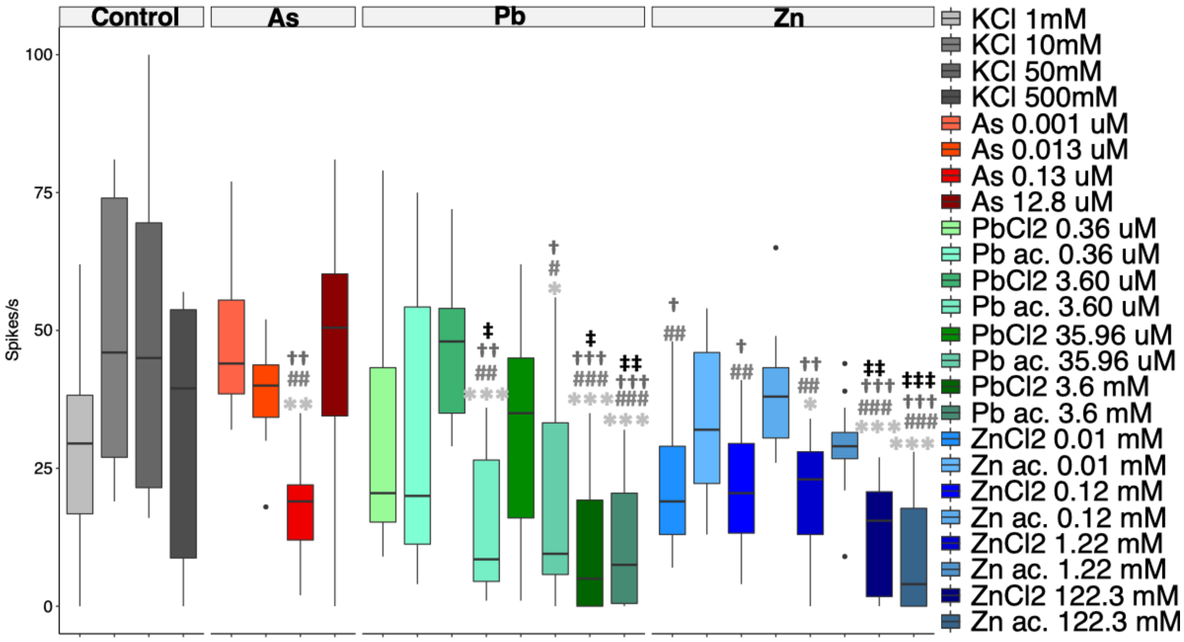


**Figure S5: Electrophysiological recordings of gustatory neurons from antennal type I sensilla.** Comparison of spike frequencies following stimulation with 30 mM sucrose containing either a common salt (KCl, grey) or metal salts (arsenic, red; lead, green; zinc, blue). P-values were obtained from GLMM, and comparisons to KCl 1mM (*), 10mM (#), 50 mM (†) and 500 mM (‡) are displayed (*$p<0.05$, **$p<0.01$, ***$p<0.001$).

**Dataset S1: Raw data (.xlsx file). 'Feeding assay – choice':** data on the feeding choice assay. Treatment, cage and hive identity, day of the experiment (1 to 3), consumption preference (g/bee/day). **'Feeding assay – no choice':** data on the feeding no-choice assay. Treatment, cage and hive identity, day of the experiment (1 to 3), consumption (g/bee/day). **'Feeding assay – survival':** data on the survival during feeding assay. Treatment, cage and hive identity, group (control, choice, no-choice), hour (1 to 72), survival (0=dead, 1=alive), number of bees. **'Antennal response assay:** data on the antennal response assay. Bee and hive identity, treatment, conditioning trial, PER (0=no response, 1=proboscis extension). **'Proboscis response assay:** data on the proboscis response assay. Bee and hive identity, treatment, conditioning trial, PER (0=no response, 1=proboscis extension). **'Survival proboscis assay':** Treatment, minute (0 to 150), survival (0=dead, 1=alive), number of bees. **'Electrophysiological recordings':** data on the electrophysiological recordings. Date, dilution (water or sucrose), type of sensilla (type I or II), bee identity, treatment, spikes frequencies.

### References


1. Monchanin C *et al.* 2021 Chronic exposure to trace lead impairs honey bee learning. *Ecotoxicol. Environ. Saf.* **212**, 112008. (doi:https://doi.org/10.1016/j.ecoenv.2021.112008)
2. Codex Alimentarius. 2015 Codex general standard for contaminants and toxins in food and feed - CODEX STAN 193-1995.
3. Codex Alimentarius. 1984 Contaminants, Joint FAO/WHO Food standards Program (Vol. XVII, 1st ed.), 163–170.
4. Pisani A, Protano G, Riccobono F. 2008 Minor and trace elements in different honey types produced in Siena County (Italy). *Food Chem.* **107**, 1553–1560. (doi:10.1016/j.foodchem.2007.09.029)
5. Bastías JM, Jambon P, Muñoz O, Manquián N, Bahamonde P, Neira M. 2013 Honey as a bioindicator of arsenic contamination due to volcanic and mining activities in Chile. *Chil. J. Agric. Res.* **73**, 18–19. (doi:10.4067/S0718-58392013000200010)
6. Czipa N, Diósi G, Phillips C, Kovács B. 2017 Examination of honeys and flowers as soil element indicators. *Environ. Monit. Assess.* **189**, 412. (doi:10.1007/s10661-017-6121-1)
7. Aggarwal I. 2017 Detection of heavy metals in honey samples using inductively coupled plasma mass spectrometry (ICP-MS). *PRAS* **1**, 1–6.
8. Terrab A, Recamales A, Gonzalezmiret M, Heredia F. 2005 Contribution to the study of avocado honeys by their mineral contents using inductively coupled plasma optical emission spectrometry. *Food Chem.* **92**, 305–309. (doi:10.1016/j.foodchem.2004.07.033)
9. Hajar EWI, Sulaiman AZB, Sakinah AMM. 2014 Assessment of heavy metals tolerance in leaves, stems and flowers of *Stevia rebaudiana* plant. *Procedia Environ. Sci.* **20**, 386–393. (doi:10.1016/j.proenv.2014.03.049)
10. Bilandžić N, Đokić M, Sedak M, Kolanović BS, Varenina I, Končurat A, Rudan N. 2011 Determination of trace elements in Croatian floral honey originating from different regions. *Food Chem.* **128**, 1160–1164. (doi:10.1016/j.foodchem.2011.04.023)
11. Al-Khalifa AS, Al-Arify IA. 1999 Physicochemical characteristics and pollen spectrum of some Saudi honeys. *Food Chem.* **67**, 21–25. (doi:10.1016/S0308-8146(99)00096-5)
12. Buldini PL, Cavalli S, Mevoli A, Sharma JL. 2001 Ion chromatographic and voltammetric determination of heavy and transition metals in honey. *Food Chem.* **73**, 487–495.
13. Maiyo WK, Kituyi JL, Mitei YJ, Kagwanja SM. 2014 Heavy metal contamination in raw honey, soil and flower samples obtained from Baringo and Keiyo Counties, Kenya. *Int. J. Emerg. Sci. Eng.* **2**, 5–9.
14. Bordean D-M, Gergen I, Harmanescu M, Rujescu CI. 2010 Mathematical model for environment contamination risk evaluation. *J. Food Agric. Environ.* **8**, 1054–1057.
15. Cozmuta A, Bretan L, Cozmuta L, Nicula C, Peter A. 2012 Lead traceability along soil-melliferous flora-bee family-apiary products chain. *J. Environ. Monit.* **14**, 1622. (doi:10.1039/c2em30084b)
16. Hussain I, Khan L. 2010 Comparative study on heavy metal contents in *Taraxacum Officinale*. *Int. J. Pharmacogn. Phytochem. Res.* **1**, 15–18.
17. Devillers J, Doré JC, Marenco M, Poirier-Duchêne F, Galand N, Viel C. 2002 Chemometrical analysis of 18 metallic and nonmetallic elements found in honeys sold in France. *J. Agric. Food Chem.* **50**, 5998–6007. (doi:10.1021/jf020497r)
18. Naggar YAA, Naiem E-SA, Seif AI, Mona MH. 2013 Honeybees and their products as a bioindicator of environmental pollution with heavy metals. *Mellifera* **13**, 10–20.
19. Przybylowski P, Wilczynska A. 2001 Honey as an environmental marker. *Food Chem.* **74**, 289–291.
20. Eskov EK, Eskova MD, Dubovik VA, Vyrodov IV. 2015 Content of heavy metals in melliferous vegetation, bee bodies, and beekeeping production. *Russ. Agric. Sci.* **41**, 396–398.

(doi:10.3103/S1068367415050079)

21. Solayman Md, Islam MdA, Paul S, Ali Y, Khalil MdI, Alam N, Gan SH. 2016 Physicochemical properties, minerals, trace elements, and heavy metals in honey of different origins: a comprehensive review. *Compr. Rev. Food Sci. Food Saf.* **15**, 219–233. (doi:10.1111/1541-4337.12182)

22. Moniruzzaman M, Chowdhury MAZ, Rahman MA, Sulaiman SA, Gan SH. 2014 Determination of mineral, trace element, and pesticide levels in honey samples originating from different regions of Malaysia compared to manuka honey. *BioMed Res. Int.* **2014**, 1–10. (doi:10.1155/2014/359890)

23. Xun E, Zhang Y, Zhao J, Guo J. 2018 Heavy metals in nectar modify behaviors of pollinators and nectar robbers: consequences for plant fitness. *Environ. Pollut.* **242**, 1166–1175. (doi:10.1016/j.envpol.2018.07.128)
